# Neovascularization and the recruitment of CD31+ cells from the bone marrow are unique under regenerative but not wound repair conditions

**DOI:** 10.1101/2022.03.02.482660

**Authors:** Kamila Bedelbaeva, Young Zhang, Azamat Azlanukov, Dmitri Gourevitch, Iossif Strehin, Phillip Messersmith, Ellen Heber-Katz

## Abstract

The long-noted observation that endostatin is a potent inhibitor of tumor vasculature but has little or no effect on wound repair or pregnancy remains an “as of yet unexplained but remarkable phenomenon”(1). However, there is another path to wound healing, epimorphic regeneration, and here we present data in mice demonstrating that endostatin is, in fact, a potent inhibitor of epimorphic regeneration. In this study, we show that a rege nerative response seen in the spontaneously regenerating MRL mouse involves CD31+ endothelial precursors that migrate from the bone marrow into the wound site and form new vessels, unlike that seen in the non-regenerating C57BL/6 mouse injury site. Furthermore, this appears to relate to the induction of HIF-1a, an inducer of regeneration (2). Inducing epimorphic regeneration in otherwise non-regenerating mice via an enhanced HIF-1a response by employing the PHD inhibitor 1,4-DPCA/hydrogel, a HIF-1a stabilizer, results in the same increased bone marrow-derived CD31+ endothelial precursor response and increased vasculogenesis. This regenerative response is completely blocked by endostatin, supporting the notion that vascularization induced during regeneration shares similarities to the tumor vasculature.

## INTRODUCTION

New blood vessels that form at wound sites or neovascularization can derive from two separable sources: The budding and extension of existing micro-capillaries into the newly forming stroma (angiogenesis) or the recruitment of bone marrow-derived endothelial precursors to wound sites (vasculogenesis) (3-6). The latter process is seen during development when new blood vessels are formed de novo (7). The relative contribution of vasculogenesis compared to angiogenesis in wound repair vs regeneration has been considered (8) but has not to date been assessed due to the fact that wound healing in adult mammals is almost exclusively via wound repair (and scarring), with the notable exception of the liver.

Hypoxia inducible factor 1a (HIF-1a) is a nodal gene which in response to hypoxia acts as a transcription factor by combining with HIF-1b or Arnt in the nucleus, binding to HIF response elements (HREs) (9) and activating over 100 genes involved in inflammation, oxygen sensing, aerobic glycolysis, and neovascularization (10-12). The latter includes the induction of VEGF, VEGFR-1/2, ANG1/2, NOS, SDF1, and PDGF to name a few (13). HIF-1a’s vital function is further evident given the vascular defects that contribute to embryonic lethality of HIF-1a mutant mice (14,15).

HIF-1a has also been shown to be a nodal gene in murine regenerative responses at multiple levels. In the regenerating MRL mouse, the most studied of a very few existing spontaneous regenerating laboratory mammals, an ear hole injury leads to the formation of a regeneration blastema (2,16). Here, a potent biphasic HIF-1a response is seen in which initial high levels of HIF-1a results in a cascade of effects including the induction of de-differentiation and stem cell markers, followed by a normalization of HIF-1a levels, increased cell proliferation and a return to homeostasis with full replacement of tissue including cartilage and new hair follicles (2,17). HIF-1a stabilization using the PHD inhibitor 1,4-DPCA delivered systemically in a hydrogel carrier (DPCA/hydrogel) also enhanced ear hole regeneration and neovascularization in a non-regenerative mouse (2,18), and importantly, recapitulated the biphasic HIF-1a response with all of the cellular dynamics of the blastema including de-differentiation and re-differentiation and the expression of the appropriate molecular markers.

In the present study, we show that epimorphic regeneration seen in ear hole closure in the MRL mouse primarily involves a strongly HIF-1a dependent vasculogenic response with CD31+ cells migrating from the bone marrow as opposed to the more HIF-1a-independent angiogenic response seen in wound repair typical of the C57BL/6 (B6) mouse. We show that the HIF-1a stabilizer 1,4-DPCA does the same thing in another non-regenerating mouse strain, Swiss Webster (SW). Secondly, we examine the effect of endostatin, an endogenous 20kDa C-terminal fragment of collagen XVIII known to be a broad spectrum vascular inhibitor (19,20), on 1,4-DPCA-induced ear hole closure, an example of epimorphic regeneration (2). With this inhibitor, ear hole closure is totally blocked. This is consistent with previously reported observations that endostatin suppresses HIF-1a expression (20,21) as we show in this study, although other reports do suggest that endostatin is HIF-independent (22). This is in contrast to studies showing that endostatin treatment of different types of cutaneous skin wounds leaves the wound repair process unaffected or even enhanced by endostatin treatment with reduced hypertrophic scar formation (23-26). Thus, responses to endostatin clearly differentiate wound repair from regeneration.

## RESULTS

### The regenerative MRL mouse shows increased CD31+ cellular accumulation and vasculogenesis after ear pinna injury and this is mimicked by 1,4-DPCA/hydrogel treatment in non-regenerative SW mice

Peak CD31 or anti-PECAM-1 expression in regenerative MRL punched ear pinnae is seen on day 7 in cells in the new growth zone of the injury site with micro-vessel/capillary formation versus little activity in non-regenerative B6 ear tissue (**Fig 1a-c**). More extensive blood vessels are found in MRL ears after co-staining with anti-caveolin (Cav1) and anti-CD31 on days 0, 7 and 15 post-injury compared to B6 (**Fig 1g-l**), supporting the idea that an increased endothelial response in the ear correlates with an increased regenerative response. Normal MRL ear tissue also shows increased levels of CAV1/CD31 (**Fig 1g**).

**Fig 1.**
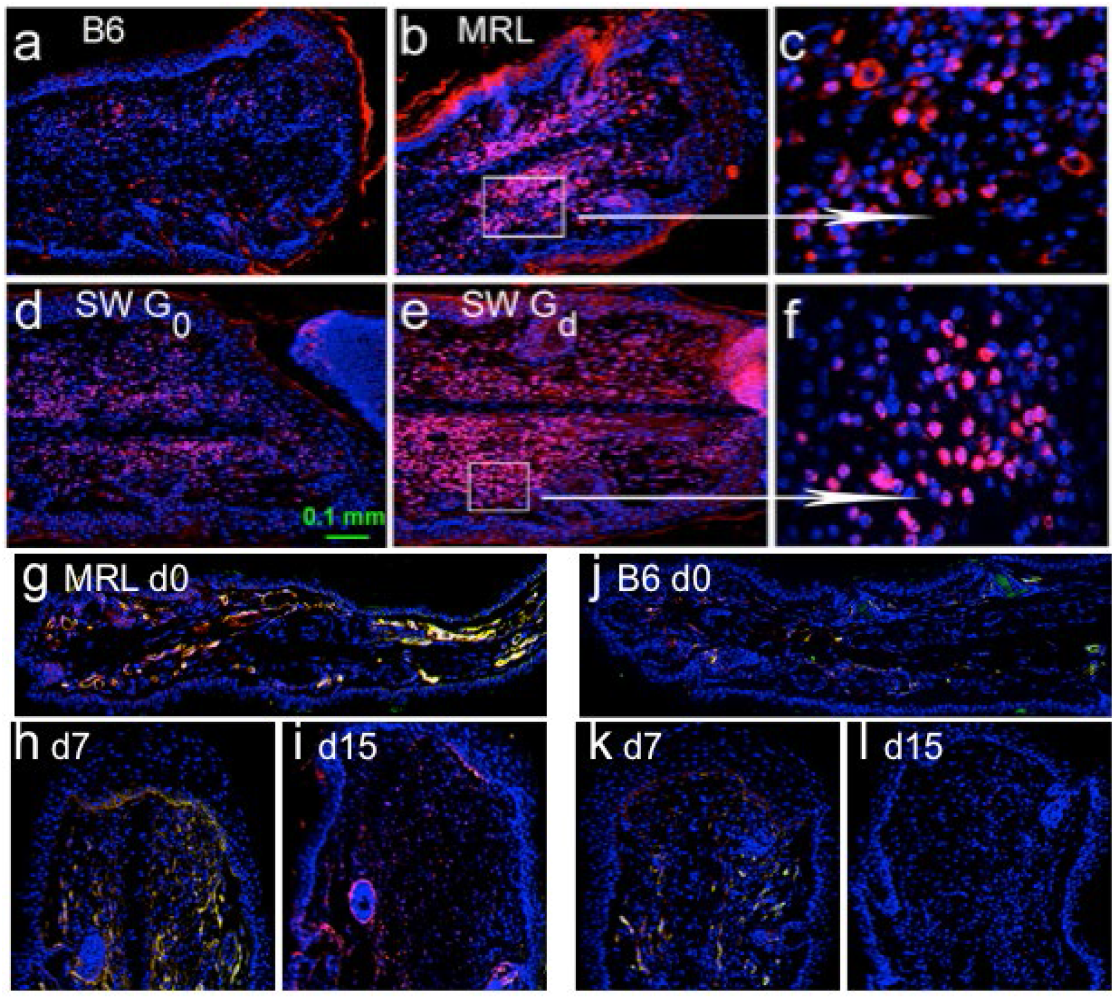
Evidence for enhanced neovascularization and vasculogenesis in regenerating ear holes: For analysis of neo-vascularization, ear tissue was immunostained with anti-CD31 (red). 1a-c compares B6 vs MRL CD31 levels da7 after injury; 1d-f compares SW G_0_ and G_d_ CD31 levels in ears da4 after injury and injection of drug/hydrogel (Gd) or hydrogel alone (G0). Here figures are at 20X except 1c,f which are at 100X magnification and represented in the white box. Next, MRL and B6 ear tissue is immunostained with both anti-Cav1 (red) and anti-CD31 (green) on days 0,7, and 15. Overlapping of those two stains produced yellow structures (1g-l). Higher double staining of blood vessels is seen in the regenerative MRL (g-i) though most of the vessels are only cav1 positive (red) on day 15. B6 shows low expression of these markers at all timepoints. NOTE: CD31 is red in 1a-f and green in 1g-l.

The regenerative MRL-type ear hole closure has been replicated in non-regenerative SW mice by treating with 1,4-DPCA/hydrogel(Gd) (**Fig 2**). As such, a similar CD31^high^ response is seen in non-regenerative SW ears post injury and DPCA/hydrogel treatment (Gd), with CD31^low^ expression in hydrogel-treated (G0) compared to the Gd-treated SW mice (**Fig 1d-f)**. This suggests that CD31^high^ expression is a regenerative phenotype as shown previously (2). The B6 and the G0-treated SW responses are typical of a wound repair response leaving a life-long open ear hole with a scarred margin (**Fig 2**).

**Fig 2.**
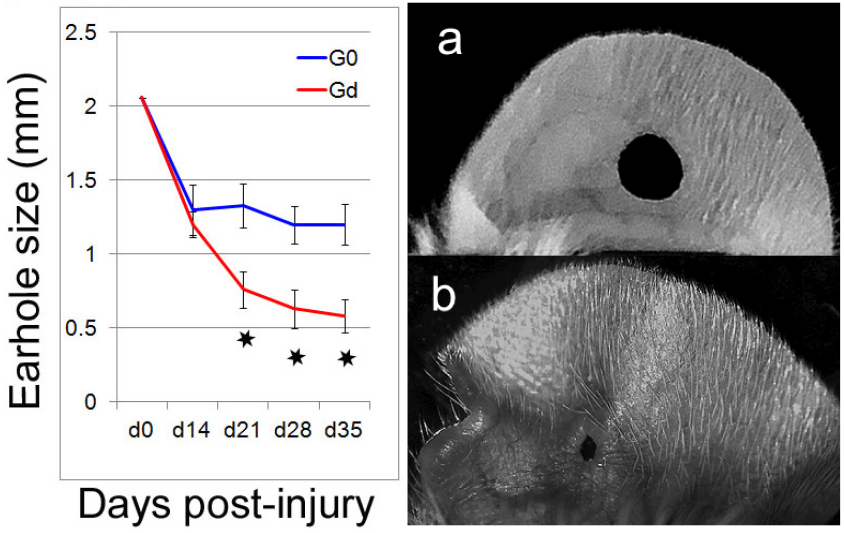
Mice injected with 1,4-DPCA/hydrogel showed enhanced ear hole closure. Non-regenerative SW mice were injected with control hydrogel (blue line, G0) or 1,4-DPCA/hydrogel (2mg/ml; red line, Gd) and earhole size in mm (hole diameter) was followed for 35 days. Mice were injected 3 times, once every five days, subcutaneously at the base of the neck, p <0.05, n=6/group. Images (right) show day 30 ear hole closure in G0(a) and Gd(b)-treated mice.

What is the source of the CD31+ cells in the spontaneously regenerating MRL mouse on day 7? Syngeneic MRL bone marrow chimeras were made by labeling MRL bone marrow cells with CFSC and injecting them into lethally (10Gy) x-irradiated MRL recipients, which are then rested for one month. This reveals that cells and vessels in the regenerating pinnae wounds are derived from injected bone marrow and are not derived from local cells. Labeled capillaries are seen on day 10 and most if not all of the vessels formed by day 15 in the MRL ear are derived from labeled bone marrow and also co-stained with anti-CD31 (**Fig 3a-d**) suggesting a vasculogenic mechanism of blood vessel growth.

**Figure 3.**
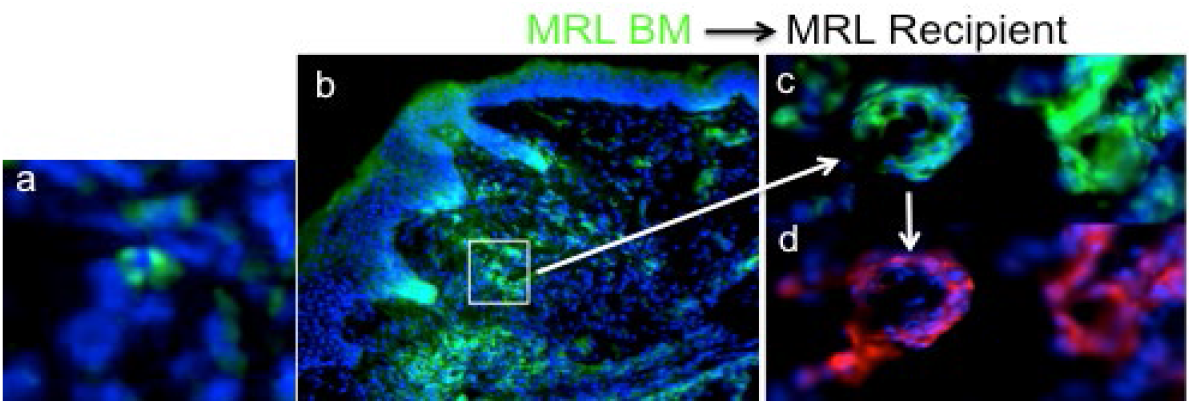
Bone marrow-derived cells in regenerative blood vessels. Chimeric MRL ear tissue from day 10 and 15 post hole-punched ears are seen. Tissue was stained with DAPI (blue) and anti-CD31 (red, Alexa 568 secondary ab) and was examined for immuno-fluorescence. Areas of green staining from CFSC-labeling of bone marrow were determined. a) At day 10 small capillaries can be seen (100x mag). b) By day 15 post-injury, large vessels have formed (40x mag). As seen in (c,d) at 100X, green stained blood vessels plus DAPI staining (c) were compared to anti-CD31-stained (red) vessels in (d). All vessels examined showed overlapping co-staining with CFSC (green) and anti-CD31 (red).

Returning to the SW model, we examined the kinetics of appearance of CD31 and von Willebrand factor (vWF) in G0- and Gd-treated injured SW ears. As seen in **Fig. 4**, Gd-treated SW mice show peak expression of CD31 on day 14 with little expression in G0-treated mouse ears. vWF is involved in blood vessel formation (27) and its expression follows CD31 and peaks on day 21.

**Fig 4.**
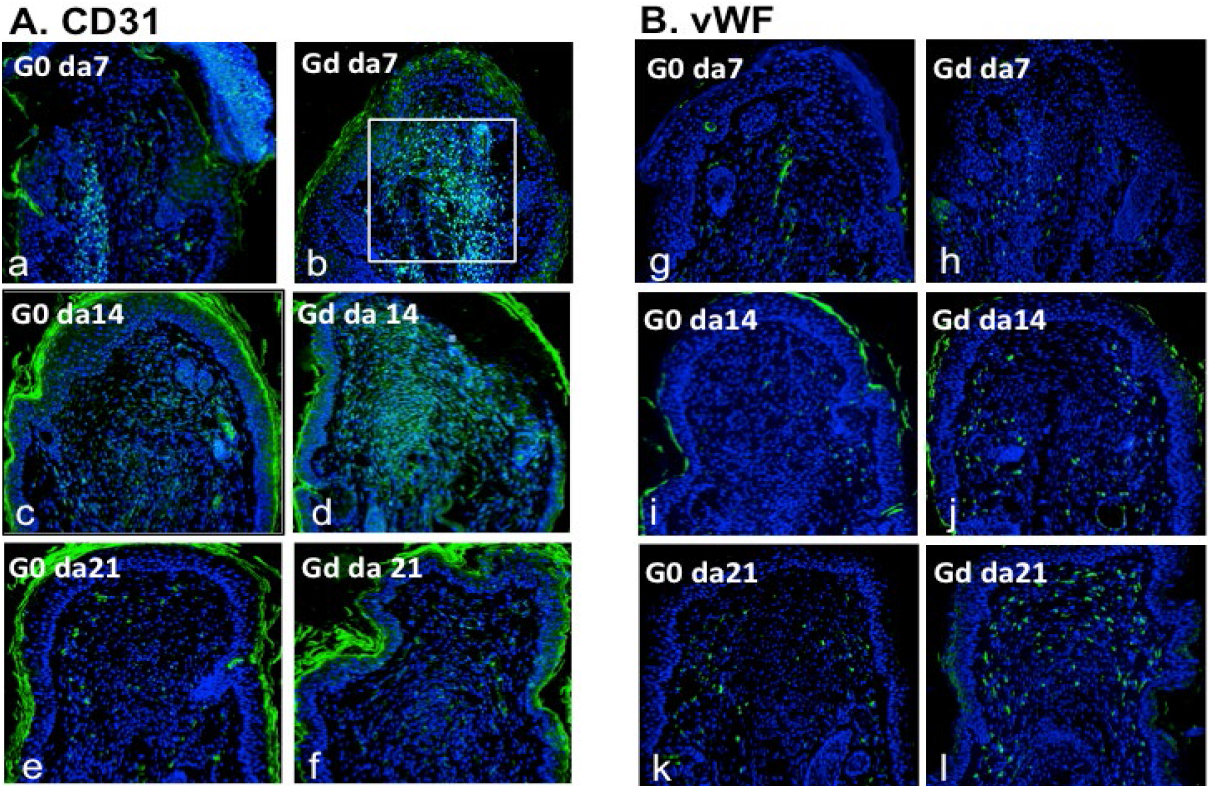
Kinetics of CD31 and vWF expression in injured ear tissue from drug-treated vs control-treated SW mice. Immuno-staining with A) anti-CD31 and B) anti-vWF shows higher staining in DPCA/hydrogel-treated SW (b,d,f and h,j,l) compared to hydrogel alone (a,c,e and g,I,k) treated ear tissue on days 7,14, and 21. Scale bar=0.1 mm. Boxed area shows blastema location. The outer layer of tissue is shed keratin with non-specific green staining.

To determine the source of the CD31-positive cells in drug-treated SW mice, we produced x-irradiation chimeric SW mice injected with CFSC (green)-labeled SW bone marrow. After one month recovery from 10 Gy x-irradiation treatment, we ear-punched these mice and began drug treatment. Ear tissue examined in G0-treated mice on day 7 and 14 display no cells that are CFSC-positive, with only a low number of CD31+ cells. On the other hand, Gd-treated mice on day 7 and day 14 show large numbers of green-labeled cells as well as blood vessels which appear after injury in the ear. These cells and capillaries are co-labeled with anti-CD31 (**Fig 5**).

**Fig 5.**
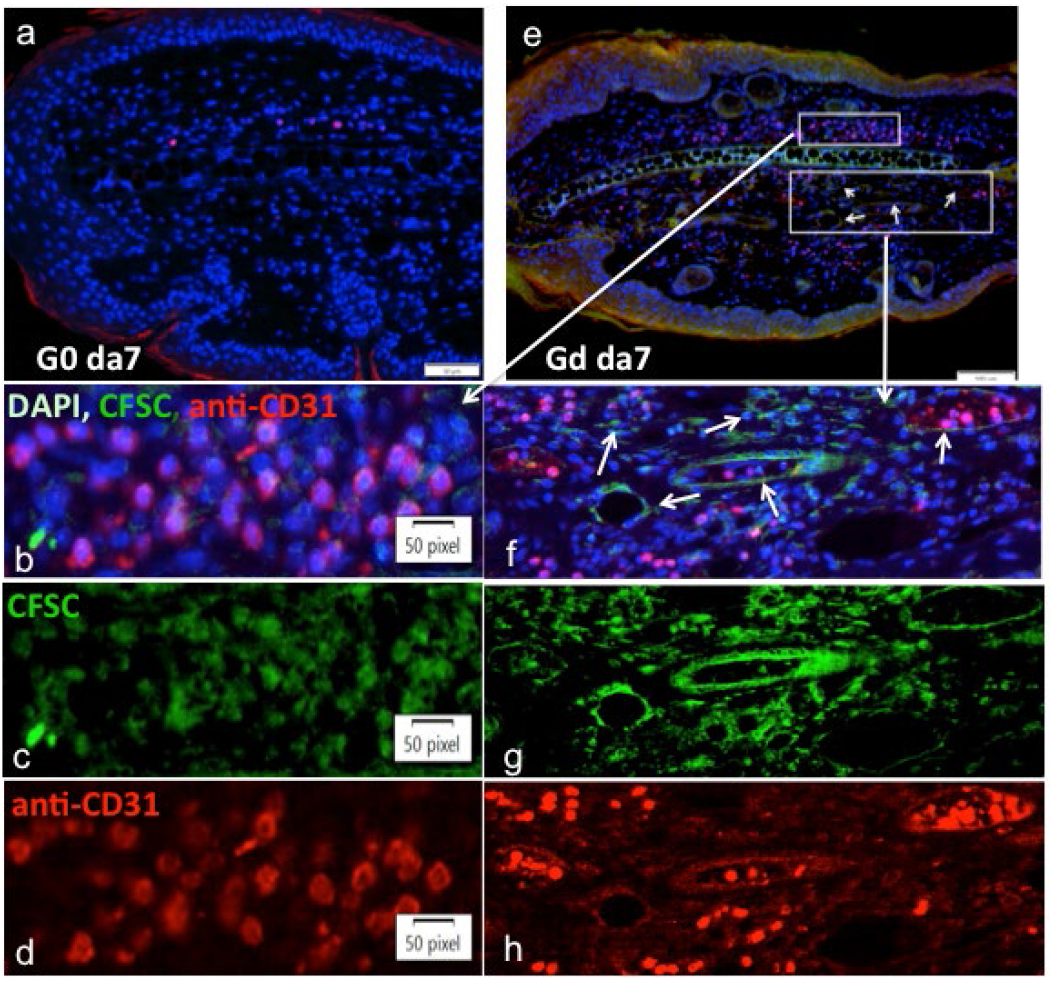
SW irradiation chimeras show BM origin of CD31+ cells suggestive of vasculogenesis. SW female bone marrow was labeled with CFSC, and injected into 10Gy x-irradiated SW female mice to generate chimeric mice. Ear tissue from these mice day 7-post hole-punching is seen. Tissue was stained with DAPI (blue) and anti-CD31 (red, Alexa 568 secondary ab) and examined for immuno-fluorescence. Ears from mice given hydrogel alone (a;G0) show no green staining and also little CD31 staining. Ears from mice given drug/hydrogel (b;Gd) show areas of staining green from CFSC-labeling of bone marrow and red from anti-CD31 staining. In the area above the cartilage (white arrow going to the left (5b-d) is shown cells co-stained with DAPI(blue), CFSC(green), and anti-CD31(red). Blood vessels found below the cartilage (boxed area with arrow to the right) can be seen to be co-stained with DAPI, CFSC, and anti-CD31 (f-h). All vessels examined show overlapping co-staining with CFSC (green) and CD31 (red). The scale bars are 50um for a, 100 um for e, and 50 pixels for b,c,d,f,g, and h.

### Blocking a regenerative response with endostatin

In an attempt to dissect and compare the vasculogenesis seen in regenerative ear hole closure vs angiogenesis in wound repair (8), we examined the effect of endostatin on G0-treated SW mice vs Gd-treated SW mice. Endostatin is known to be a neovascular inhibitor but has long been noted not to affect wound repair. What, however, would be its effect on vasculogenesis in the context of regeneration? And, since 1,4-DPCA/hydrogel upregulates HIF-1a expression, a requirement for regeneration (2), how does the effect of endostatin compare to *siHif1a*?

As seen in **Fig 6**, endostatin, like si*Hif* (2), showed almost complete blockade of regenerative ear hole closure in these mice, totally different from its lack of effect on wound repair (24). Endostatin and si*Hif* had differing downstream effects, however, on the expression of molecular markers of angiogenesis, the stem cell state, tissue remodeling and inflammation, including HIF target genes allowing an ordering of some processes central to regeneration (2).

**Fig 6.**
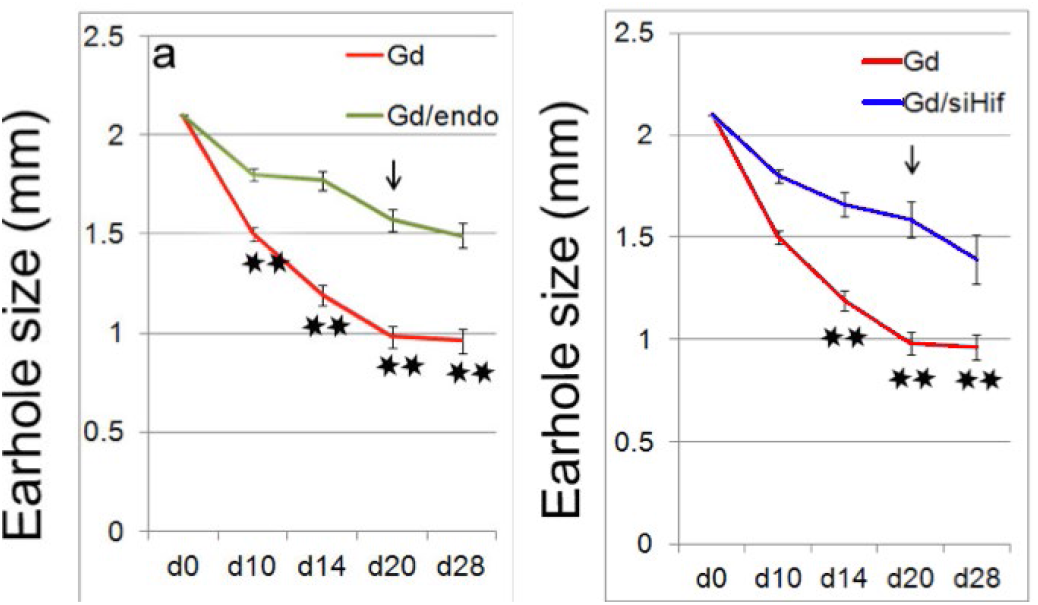
The effect of endostatin or si*Hif1a* on ear hole closure of Swiss Webster mice treated with 1,4-DPCA/hydrogel. Kinetics of ear hole closure with and without endostatin (left) or *siHif1a* (right) treatment is seen. All time points examined showed highly significant differences (**P<0.01). Both endostatin and si*Hif* were given every other day up to 20 days (arrow indicates end of treatment).

Immunohistochemistry (IHC) was carried out on ear tissue from SW G0- and Gd-treated mice, and from Gd-treated mice which then received either endostatin or si*Hif1a* (**Fig 7a-p**). These results were quantified using ImagePro v4.0 and the area was expressed in square microns (**Fig 7q)**. Note, all Gd-treated mice are committed to the regeneration pathway.

**Fig. 7.**
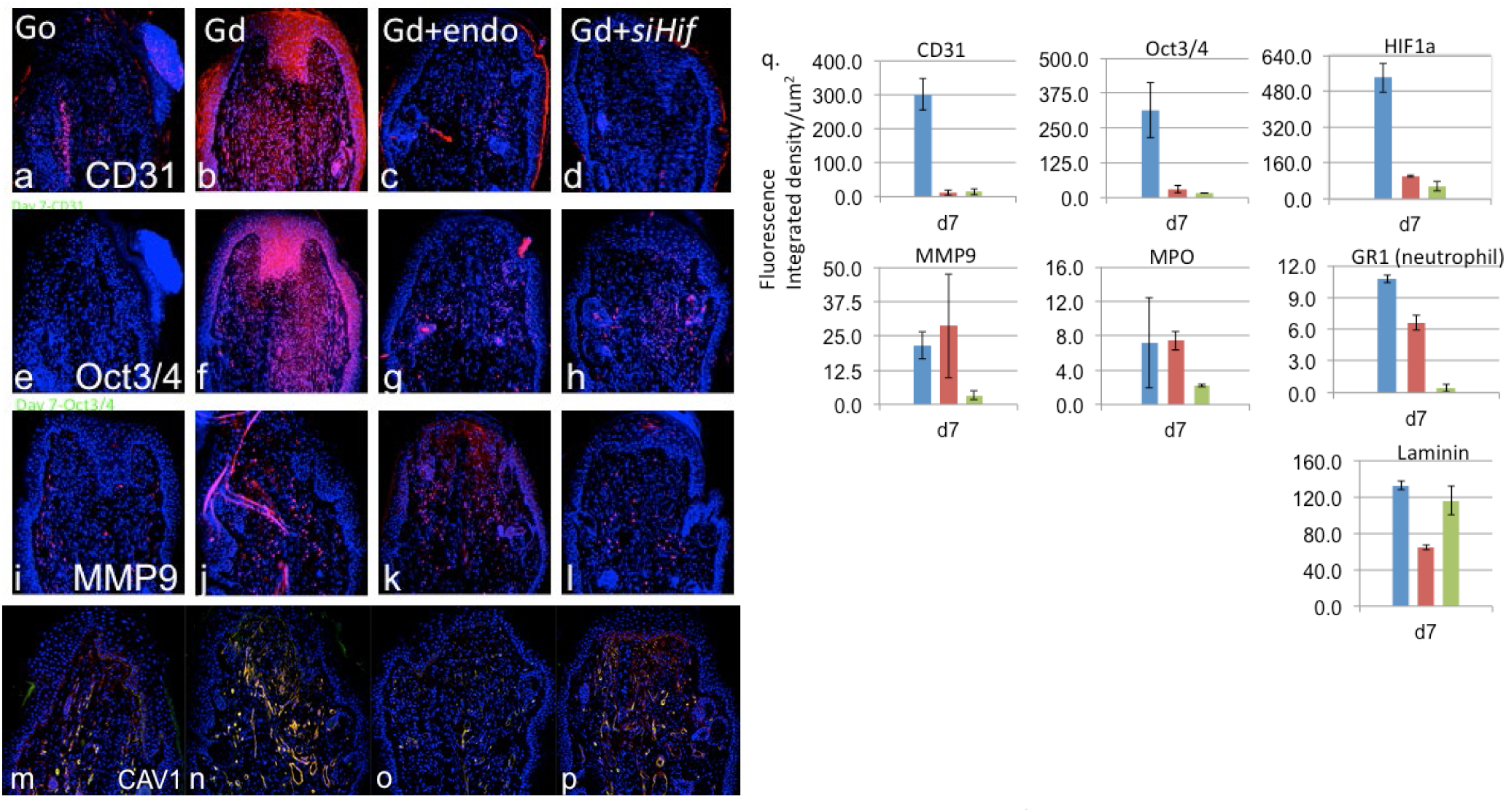
The immunostaining of ear tissue from SW mice injected with hydrogel alone (G0; 7a,e,i & m), with 1,4-DPCA/hydrogel (Gd; 7b,f,j & n), with Gd + endostatin (7c,g,k & o) or with Gd + si*Hif1a* (7d,h,l & p). The molecules examined include CD31 (7a-d), Oct3/4 (7e-h), MMP9 (7i-l), and cav1+CD31 (7m-p). The secondary ab were Alexa (red) and DAPI (blue). (q) Quantitation of IHC analysis is seen for G_d_ (blue bars), G_d_ + endostatin (red bars), and G_d_ + si*Hif1a* (green bars) injected mouse-derived tissue. Data are shown as the fluorescence integrated density/um^2^ mean +/− SE from samples (n=5) per treatment group. (Cav1-red goat anti Rabbit Alexa Fluor 568 nm; CD31- donkey anti goat green Alexa Fluor 488 nm.

On day 7-post injury plus treatment, three different phenotypes are seen. In the first phenotype, both endostatin and si*Hif* almost completely blocked the expression of CD31, an endothelial cell progenitor marker, HIF-1a, and OCT3/4, an embryonic stem cell transcription factor and marker of undifferentiated cells. In the second phenotype, the matrix metalloproteinase and remodeling molecule, MMP-9, as well as the pro-inflammatory markers MPO and GR-1 (a neutrophil marker) were inhibited by si*Hif* but to a much lesser degree or not at all with endostatin. In the third phenotype, a greater degree of inhibition was seen with endostatin compared to si*Hif*. This included mature blood vessel markers laminin and caveolin (not graphed, but histology is shown in Fig 7m-p).

## DISCUSSION

In the typical wound repair site, the wound experiences a hypoxic response which includes the enhancement of a functional vascular network allowing increased oxygen flow by increasing the number of both red blood cells and blood vessels. This is achieved by the reduction in prolyl hydroxylase (PHD) activity, the stabilization of HIF-a proteins, the formation of HIFa/b transcription factors in the nucleus, and the activation of their downstream targets including VEGF, PDGF, bFGF, MCP-1, HMOX-1 and erythropoietin (EPO) increasing blood cell numbers. Besides these targets, HIF-1a is a key molecule that is involved in a multitude of functions of the cell, from regulating O2 levels and erythropoesis to aerobic glycolysis, cell migration, and inflammation (9-13). This process results in a scar. On the other hand, in scarless regenerative healing, there is a massive increase in HIF-1a for 10-14 days leading to quantitative and qualitative differences. As we showed previously (2), blocking the HIF-1a response abrogates regeneration.

We had noted that the regenerative adult MRL mouse uses aerobic glycolysis as its normal metabolic state (28) and it was thus not surprising to find that HIF-1a protein levels were unusually high in this mouse during early wound healing, as demonstrated by ear hole closure, including blastemal expansion and de-differentiation (2). Regenerative healing was completely blocked by si*Hif1a.* This response was replicated in non-regenerative mice using the HIF-1a-stabilizing drug 1,4-DPCA incorporated into a hydrogel (2,18) and given subcutaneously distal to the wound. Thus, the “MRL regenerative phenotype” could be replicated in every aspect examined and si*Hif* could abrogate this response as well. This type of healing has a biphasic curve in which HIF-1a levels rise for approximately 10-14 days and is accompanied by enhanced tissue remodeling, increased MMP levels, a de-differentiated cellular signature, increased glycolytic enzymes, increased components of inflammatory and enhanced cell migration and proliferation leading to ear hole closure. In the second half of the biphasic curve, HIF-1a levels fall with increased re-differentiation to mature cartilage and hair follicles. In the current studies, SW non-regenerative mice were given 1,4-DPCA in hydrogel, described previously, to explore its effect on neovascularization, a major regulatory target of HIF-1a.

### Neovascularization

The need for new blood vessels during wound healing is well described (3,6,8,29). It has been shown that such vessels are generated by two mechanisms. The first, angiogenesis, is fueled by endothelial cells derived from pre-existing vessels, which migrate, proliferate, and form sproutings from the local mature vessels. The second, vasculogenesis, is the development of blood vessels from CD31+ endothelial precursor cells released from the bone marrow which then migrate through the blood to wound sites and is generally used during development to form blood vessels de novo.

The massive appearance of CD-31+ cells in regenerative tissue wounds as opposed to the lack of these cells in non-regenerative wounds along with the bone marrow source of those CD31+ cells strongly points to the more developmental process of vasculogenesis during regeneration. Furthermore, the blocking effect of endostatin in the regenerative response supports this. It has been proposed that increased angiogenesis leads to increased scarring during wound repair (29) that is seen in the non-regenerating non-closing ear hole wounds. Vasculogenesis, on the other hand, seen in fetal wounding correlates with scarless healing. Thus the idea that vasculogenesis in adult regeneration could lead to non-scarred healing is supported by what is found in the MRL as well as in Swiss Webster mice treated with 1,4-DPCA/hydrogel.

A role for neovascularization in ear hole injuries has been noted in previous studies. The AGF (angiopoietin-related growth factor) transgenic mouse with increased vascular and epithelial proliferation showed an enhanced ear hole closure response in 28 days, similar to the MRL (16,30). Angiopoietin (angpt1) showed different effects. The angpt KO mouse displayed positive ear hole closure (31) which was thought to be due to an increased angiogenic and fibrotic but not a regenerative response. Ear hole closure took 60 instead of the usual 30 days and never regrew cartilage or hair follicles, unlike the MRL mouse (16). In the adult, Angpt balances enhanced levels of TgFb, Vegfa, and Angpt2 and inhibits angiogenesis and fibrosis. In a third study, recombinant COMP-ang1 was injected into ear-punched non-regenerative mice with some closure seen but never reaching a healing phenotype by day 28 (32). The only regenerative response was achieved with AGF, though cartilage growth and hair follicle growth was not examined.

In this study, we found that endostatin, a broad spectrum neovascular inhibitor, could completely block ear hole closure in regeneration-competent mice. We compared endostatin to si*Hif* for their ability to block 1,4 DPCA-induced hole closure and found that all of the regenerative phenotypes described previously (2) were affected by *siHif* and/or endostatin, some similar and some the opposite allowing us to generate preliminary associations of effector functions.

The first phenotypic group includes three molecules, HIF-1a, CD31, and OCT3/4, which are inhibited equally by si*Hif* and endostatin. HIF-1a has been previously shown to be down-regulated by endostatin (19-21) so that many of the inhibitory results could be due to inhibition of HIF-1a or by other endostatin-specific interactions. CD31 is a HIF-1a target and is expressed by endothelial cells, blood vessels, and cells migrating from the bone marrow. OCT3/4, a marker of de-differentiation, though not a direct HIF target, is activated by HIF-1a (2). Furthermore, the de-differentiation response, seen in the MRL and induced by HIF-1a is likely due to chromatin remodeling (33,34). One possible explanation for endostatin inhibitory effects may be completely due to HIF inhibition. However, PHD inhibitors such as 1,4-DPCA, besides stabilizing HIF-1a can also enhance NFKB activation (35,36) and NFKB can regulate the recruitment of HDACs or HATs (37) and lead to chromatin remodeling and expression of OCT3/4.

The second phenotypic group includes inflammatory components and are completely blocked by *siHif* but not by endostatin. This includes myeloperoxidase (MPO), a heme protein made during myeloid differentiation and a major component of neutrophil granules, MMP-9, a protease which has a zinc-binding domain and is involve in angiogenesis and tissue remodeling, and anti-GR1, a marker of neutrophils, seen in the MRL and induced by 1,4-DPCA/hydrogel. This is puzzling because endostatin has been shown to block inflammatory responses in rheumatoid arthritis (RA) (38) as well as NFkB, the main inflammatory nodal gene/transcriptional factor (35). It may be related to responses specifically in the ear pinna.

The third group, is mainly affected by endostatin. This is true for the molecules laminin and caveolin, the former being a major basement membrane component (39) and the latter functioning as a blood flow mechanosensor (40). These two molecules are expressed later in vessel development and at high levels in mature blood vessels. They are not affected by siHIF and are not HIF-1a target genes. As previously discussed (2,18), the HIF-1a response is biphasic, the second half being the lowering of HIF levels. High endostatin levels may both turn off HIF-1a as well as down-regulate vessel growth and lead to maturation.

Collectively, these data suggest that it is vasculogenesis that is being blocked completely by endostatin and *siHIF1a*, due to inhibition of the migration and homing of bone marrow-derived endothelial precursors to the regenerating ear wound. In contrast, the angiogeneic process of wound repair of cutaneous injuries are insensitive to endostatin. It should be noted that the SW control (G0) shows a wound repair response where the ear punch wound margin forms a scar and an open ear hole remains.

There is yet another interesting association. There is a paradoxical effect of endostatin in tumors versus wound repair and these new results may suggest the beginning of a resolution (1). While wound healing by the process of regeneration blastema formation clearly does not constitute a tumor, it has often been noted that tumors and regeneration blastemas do share some resemblance (41-46). Regeneration blastemas are seen to employ an endostatin-sensitive vasculogenic mechanism like tumors on the one-hand and are in contrast with the endostatin-resistant wound repair process which employs angiogenesis.

Finally, the mobilization of bone marrow progenitor cells in adults should enhance vascular regeneration and many homeostatic processes, recovery from injury, ischemic diseases in muscle and cardiac tissue and in age-related diseases (47). It has also most recently become important to consider the vascularization of implanted engineered tissue in vivo (48,49). Having a method to enhance such mobilization or increase migration into wound sites, besides the use of cytokines, chemokines, and growth factors (50,51), is of interest (2,52).

## Acknowledgements and Funding

We appreciate the generous support of these studies from grants from the NIDCR, NIH, from DARPA, DOD, and from the Leila Y and Harold G Mathers Foundation and the FM Kirby Foundation.

## Author Contributions

EHK designed the studies, analyzed the data and interpreted the results and wrote the paper. YZ, AA, and LC carried out the animal experiments. KB carried out the immunohistochemistry and cell culture, DG carried out the histology, data analysis, and figure preparation.

## Competing Interests

No consulting positions; US patent applications

## MATERIALS AND METHODS

### Animals and in vivo procedures

MRL/MpJ female mice were obtained from The Jackson Laboratory; C57BL/6 female mice were obtained from Taconic Laboratories; Swiss Webster female mice were obtained from Charles River. All mice were used at 8-10 weeks of age in all experiments under standard conditions at the Wistar Institute and Lankenau Institute for Medical Research Animal facilities and the protocols were in accordance with the National Institutes of Health *Guide for the Care and Use of Laboratory Animals*. Through-and-through ear hole punches were carried out as previously described and each earhole diameter was measured on the long axis and perpendicular axis of the ear pinna.

Swiss Webster mice were injected subcutaneously at the base of the neck with 100 ml of 10% (w/v) 1:1 (w/w) ratio of P8Cys (with or without 1,4-DPCA drug crystal) to P8NHS hydrogel prepared in PBS (Fig. 1 and 2) and Supplementary Materials). Each component was kept cold and mixed just before injection. At different time points, mice were euthanized and tissues were removed for protein and RNA analysis.

Bone marrow chimeras were generated by isolating bone marrow from MRL or Swiss Webster female mice, incubating and staining with CFSE, and then reconstituting lethally x-irradiated (1000R or 10Gy) syngeneic mice with bone marrow injected intravenously through the tail vein (bone marrow from 1 mouse (10^8 cells) into 3 mouse recipients.) After 1 month, mice were ear punched and followed for CFSC and CD31+ responses in the ear.

### Inhibitor treatment in vivo

#### SiHif1a

si*Hif1a* was ordered from Qiagen and siRNA Mm_HIF1a_3 (S100193025) was selected as the best inhibitor for Swiss Webster. In vivo, this siRNA was used for HIF-1a inhibition. Si*Hif* was used at 75mg/kg and mixed with Jetpei (Polyplus, Genycell) following the manufacturer’s instructions. The mice were injected subcutaneously every 48 hours up to 20 days.

#### Endostatin

Endostatin was obtained from PROSPEC-Tany TechnoGene Ltd (Israel). Mice were injected subcutaneously at a dose of 20 ug/mouse in 100 ul PBS every 48 hours up to 20 days.

### Immunohistochemistry

The methods used were performed as previously described (53). Tissue from normal ears were fixed with Prefer fixative (the active ingredient is glyoxal) (Anatech) overnight and then washed in H20. Tissue was embedded in paraffin and 5-μm thick sections cut. Before staining, slides were dewaxed in xylene and rehydrated. Antigen retrieval was performed by autoclaving for 20 min in 10 mM Sodium Citrate, pH 6.0. Tissue sections were then treated with 3% H2O2 and nonspecific binding was blocked with 4% BSA (A7906; Sigma) for 1 h. The primary antibodies and matched secondary antibodies used for IHC were shown in Table S1. For immunocytochemistry staining, primary ear skin fibroblasts were grown on coverslips in DMEM with 10% FBS at 37 °C in a humidified 5% CO2 incubator. The coverslips were rinsed with 1× PBS, the cells were fixed in cold methanol (−20 °C) for 10 min, rinsed with 1× PBS, treated with 0.1% Triton-X100, and then incubated with the appropriate primary and secondary antibodies (Table S1). Photomicrographs were produced using the fluorescent microscope (Olympus AX70).

For quantitation of IHC signal, the method used was previously described (54). Briefly, we used ImagePro v4.0 for image analysis by selecting positive staining from multiple areas in the sections. The number of “positive staining” or “non-black” pixels after conversion to gray scale was determined. The area was expressed in square microns and the final data were expressed as IHC staining signal per square micron. The mean of 2–6 samples were plotted and standard errors calculated.

### Statistical analysis

The data represent pooled samples for analysis of hole diameter for healing studies and tissue analysis (n). Student’s t test was carried out to compare differences of means from independent samples between two groups. All error bars shown on the graphs represent SEs. Statistical analyses were done using Microsoft Excel 2010.

### Hydrogel and Drug Preparation as described previously. (15,55,56)

**Supplemental Table S1.**
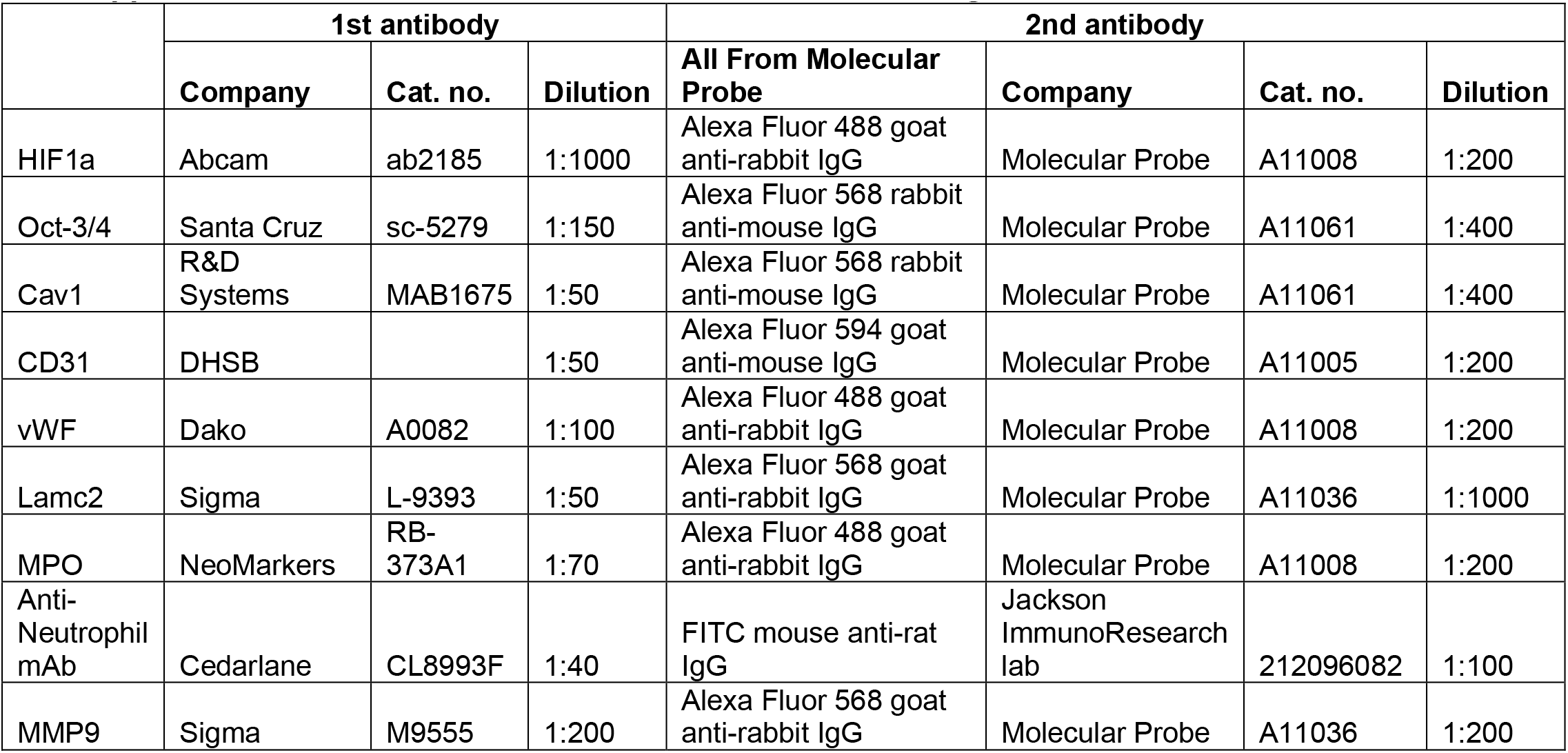
Antibodies used for Immunostaining.

## Notes

### Competing Interest Statement

The authors have declared no competing interest.

